# Urinary Proteome Characterization of Stroke-Prone Spontaneously Hypertensive Rats

**DOI:** 10.1101/2024.11.12.623308

**Authors:** Wenshu Meng, Youhe Gao

## Abstract

**Background:** Hypertension is a multifactorial, complex disease related to genetic and environmental factors and has become the most serious public health problem. This study aimed to explore the changes of hypertension based on urinary proteome.

**Methods:** The stroke-prone spontaneously hypertensive rats (SHRSPs) model was used to examined urinary proteome changes during hypertension development. Urine proteome at months 1, 4, 8, 10, 12, and 14 was profiled using liquid chromatography coupled with tandem mass spectrometry (LC-MS/MS). Considered the degree of disease progression may differ, each rat was compared before and after hypertension developed to screen for differential proteins.

**Results:** The differential proteins in each rat can be enriched into some important biological processes and pathways associated with hypertension, such as regulation of systemic arterial blood pressure by renin-angiotensin, renin-angiotensin signaling, response to glucocorticoid and glucocorticoid receptor signaling, calcium transport I, aldosterone adipocyte signaling pathway, apelin adipocyte signaling pathway and oxidative stress response. The biological processes and pathways enriched at the same time point in the progression of hypertension differed significantly among different rat individuals.

**Conclusions:** This study showed that the changes of hypertension can be reflected in urine proteins. The urine proteomics has the potential to be used to study the mechanisms of hypertension, discovery new drug targets, and provide personalized antihypertensive treatment strategies.

## Introduction

Hypertension is a clinical syndrome primarily characterized by elevated systemic arterial blood pressure in the systemic circulation. Over the past thirty years, the number of individuals with hypertension globally has reached 1.3 billion, making it a major public health problem(1, 2). Hypertension is directly associated with over 8.5 million deaths worldwide each year, and it is a leading risk factor for stroke, ischemic heart disease, and kidney disorders(3-5). Effective treatment and control of hypertension will significantly reduce the incidence and mortality rates associated with diseases related to elevated blood pressure(6). The main mechanisms underlying hypertension include the sympathetic nervous system, the renal and adrenal function, the endothelium, and the insulin resistance(7, 8).

Despite of availability and use of various antihypertensive agents, the prevalence of hypertension and related death toll caused by its complications show no signs of slowing down, indicating that important pathophysiological mechanisms of hypertension are still not well understood(9). On the one hand, due to the existence of individual differences, the therapeutic effect of antihypertensive drugs is unstable, which also brings challenges to the choice of clinical antihypertensive drugs; on the other hand, there is a lack of research on the pathologic mechanisms at different stages in the progression of hypertension. Therefore, new methods are urgently needed to elucidate the pathogenesis of hypertension and possible personalized treatment strategies to improve the rate of blood pressure control.

Proteomics can study endogenous changes and the impact of the external environment on pathophysiological processes as a whole, serving as an effective tool for discovering disease biomarkers, exploring mechanisms of disease, and screening potential drug therapy targets(10). Without the homeostatic regulation, urine can sensitively reflect changes in the body at an early stage (11). Urine is a promising resource for biomarkers research and urine proteomics has been applied to the clinical research of multiple diseases, such as gastric cancer(12), lung cancer(13), pediatric medulloblastoma(14), and Parkinson’s disease(15). Recent studies have shown that the urinary proteome has potential for application in the field of cardiovascular disease (16). For example, by comparing urinary samples from patients with carotid artery stenosis (CAD) to healthy controls, urinary biomarkers have been identified for early screening and risk stratification of CAD (17). In addition, a urinary proteomic biomarker model has been developed to assist in the diagnosis of acute stroke in those with mild symptoms(18). As a completely non-invasive sample, urine sample is very suitable for the collection at various stages of disease progression to dynamic monitoring the disease. However, the number of proteomic studies for primary hypertension is relatively limited. In this research, we aimed to determine whether the urine proteome can reflect changes associated with hypertension.

The urine proteome is easily affected by external factors, such as age, diet, exercise, sex, medication, and daily rhythms(19). Animal models can minimize the impact of many uncertain factors, thereby establishing a direct relationship between the disease and corresponding urinary changes. Stroke-prone spontaneously hypertensive rats (SHRSP) model is a valuable animal model to investigate human hypertension. In this model, the elevation of blood pressure in rats is determined by polygenic inheritance, which is similar to human hypertension (20-23).

In this study, we employed the SHRSP model to simulate the occurrence and progression of hypertension, and analyzed urine samples from six time points during the development of hypertension (1, 4, 8, 10, 12, and 14 months) using liquid chromatography coupled with tandem mass spectrometry (LC-MS/MS) with the data-independent acquisition (DIA) approach. This study screened differential proteins by comparing before and after time points to dynamically explore the pathological mechanisms of hypertension. The workflow of this study is presented in **Figure 1**.

**Fig. 1.**
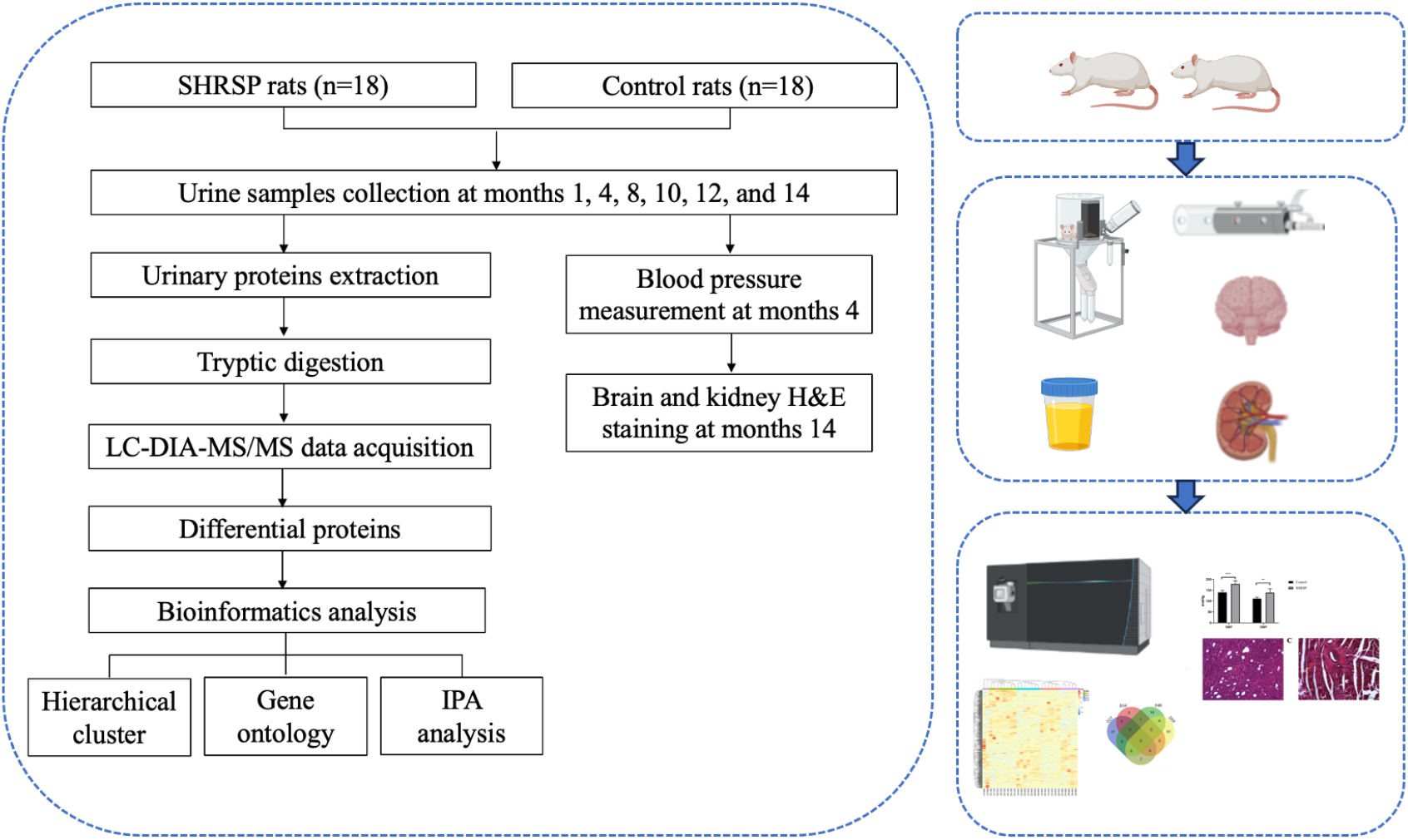
Workflow of the study of urinary proteomics changes in SHRSP rats.

## Materials & Methods

### Experimental animals and model establishment

Male SHRSPs (n=18, 140 ± 20 g) and male Wistar rats aged 4 weeks (n=6, 140 ± 20 g) were purchased from Beijing Vital River Laboratory Animal Technology Co., Ltd. Animals were fed a standard laboratory diet and housed under controlled indoor temperature (21 ± 2 °C), humidity (65-70%) and 12 h/12 h light-dark cycle conditions. The study was approved by Peking Union Medical College (Approval ID: ACUC-A02-2014-007) and performed according to the guidelines developed by the Institutional Animal Care and Use Committee.

### Urine collection and sample preparation

Urine samples were collected from the experimental group in months 1, 2, 4, 8, 10, 12 and 14. Animals were individually placed in metabolic cages for 10 h to collect urine samples without any treatment (from 8 a.m. to 6 p.m.). After collection, the urine samples were quickly stored at -80 °C. The urine samples were centrifuged at 12,000 g for 40 min at 4 °C to remove cell debris. The supernatants were precipitated with three volumes of ethanol at -20 °C overnight and then centrifuged at 12,000 g for 30 min. The pellet was resuspended in lysis buffer (8 mol/L urea, 2 mol/L thiourea, 50 mmol/L Tris, and 25 mmol/L DTT). The protein concentration of the urine samples was measured using the Bradford assay.

### Measurement of blood pressure

At the age of 4 months, a noninvasive blood pressure measuring instrument was used to measure the blood pressure at the tail artery as previously described (24). Briefly, rats were fixed on a fixation frame at room temperature until the body temperature increased to 39 °C. The proximal tail was attached to the computer via a dynamic signal acquisition system. The mean BP was obtained from three consecutive readings at 3 min intervals.

### Histopathology

At the age of 14 months, 3 rats were subjected to intracardiac perfusion. Whole-body blood was quickly flushed with 0.9% normal saline and tissues were fixed with 4% paraformaldehyde. The brain and kidney were removed and preserved in 4% paraformaldehyde. Then, the samples were embedded in paraffin, sectioned, and evaluated with hematoxylin and eosin (H&E) staining.

### Protein digestion

One hundred micrograms of urinary proteins from each sample were digested with trypsin (Trypsin Gold, Mass Spec Grade, Promega, Fitchburg, WI, USA) using filter-aided sample preparation (FASP) methods as previously described (25). These peptide mixtures were desalted\ using Oasis HLB cartridges (Waters, Milford, MA) and dried by vacuum evaporation (Thermo Fisher Scientific, Bremen, Germany). The digested peptides (n=36) were redissolved in 0.1% formic acid to a concentration of 0.5 µg/µL, and 1 µg of peptides from each sample was analyzed using LC–MS/MS in DIA mode.

### Reverse-phase fractionation spin column separation

A pooled sample was generated from equal volumes of digested peptides from each sample. A total of 108 µg of pooled peptides was separated using a high-pH reversed-phase peptide fractionation kit (Thermo Pierce, Waltham, MA, USA) according to the manufacturer’s instructions. A step gradient of increasing acetonitrile concentrations (5, 7.5, 10, 12.5, 15, 17.5, 20 and 50% acetonitrile) was added to the columns to elute peptides, and ten different fractions (including the flow-through fraction, wash fraction, and eight step gradient sample fractions) of each sample were collected and dried by vacuum evaporation. The ten fractions were then resuspended in 20 µl of 0.1% formic acid, and 1 µg of peptides from each fraction was used for LC–MS/MS analysis in DDA mode.

### LC–MS/MS analysis

Mass spectrometry acquisition and analysis were performed using an EASY-nLC 1200 chromatography system (Thermo Fisher Scientific) and an Orbitrap Fusion Lumos Tribrid mass spectrometer (Thermo Fisher Scientific). The iRT reagent (Biognosys, Switzerland) was spiked at a concentration of 1:10 v/v into all urinary samples for calibration of the retention time of the extracted peptide peaks. The peptide samples were loaded on a trap column (75 µm × 2 cm, 3 µm, C18, 100 Å) and a reverse-phase analysis column (75 µm × 25 cm, 2 µm, C18, 100 Å). The elution gradient was 4%-35% buffer B (0.1% formic acid in 80% acetonitrile) at a flow rate of 400 nL/min for 90 min. One microgram of each fraction from the spin column was analyzed in DDA mode to generate the spectral library. The parameters were set as follows: the full scan was acquired from 350 to 1550 m/z with a resolution of 120,000, and the MS/MS scan was performed with a resolution of 30,000 in the Orbitrap; the higher-energy collisional dissociation (HCD) energy was set to 30%; the automatic control (AGC) target was set to 5.0e4; and the maximum injection time was set to 45 ms.

One microgram of each sample was analyzed in DIA mode. The variable isolation window of the DIA method with 36 windows was set for DIA. The parameters were set as follows: the full scan was acquired from 350 to 1500 m/z with a resolution of 60,000; the MS/MS scan was acquired from 200 to 2000 m/z with a resolution of 30,000; the HCD energy was set to 32%; the AGC target was set to 1.0e6; and the maximum injection time was set to 100 ms.

### Data analysis

The DDA data from the ten fractions were processed using Proteome Discoverer software (version 2.1, Thermo Scientific) and searched against the Swiss-Prot rat database (released in 2017, including 7992 sequences) appended with the iRT peptide sequence. The search parameters were set as follows: two missed trypsin cleavage sites were allowed, the parent ion mass tolerances were set to 10 ppm, the fragment ion mass tolerances were set to 0.02 Da, the carbamidomethyl of cysteine was set as a fixed modification, and the oxidation of methionine was set as a variable modification. The false discovery rate (FDR) of proteins was less than 1%. A total of 862 protein groups, 4570 peptide groups and 43277 peptide spectrum matches were identified. The search results were used to set the variable windows for DIA.

Ten DDA raw files were processed using Spectronaut Pulsar X (Biognosys, Switzerland) with the default parameters to generate the spectral library. Then, 36 raw DIA files for each sample were processed using Spectronaut Pulsar X with the default settings. The results were filtered based on a q value cutoff of 0.01. The peptide intensity was based on the peak areas of the respective fragment ions for MS2, and the protein intensity was calculated by summing the intensities of their respective peptides.

### Statistical analysis

The k-nearest neighbor (K-NN) method was used to fill the missing values of protein abundance (26). As the degree of disease progression may differ, each rat was compared at time points before and after hypertension development to screen for differentially expressed proteins. The differential proteins were screened based on the following criteria: proteins with at least two unique peptides, fold change ≥ 1.5 or ≤ 0.67, and P < 0.05 in a two-sided unpaired t test. Group differences resulting in P < 0.05 were considered statistically significant.

### Functional annotation of the differential proteins

The Database for Annotation, Visualization and Integrated Discovery (DAVID) was used to perform the functional annotation of the differential proteins and included biological processes, cellular components and molecular functions (27). The canonical pathways were analyzed with IPA (Ingenuity Systems, Mountain View, CA, USA) software. All enriched terms had a threshold value of P < 0.05.

## Results

### Characterization of SHRSP rats

The systolic blood pressure (SBP) and the diastolic blood pressure (DBP) measured by the tail-cuff method showed significant difference between the SHRSP and the control rats at Month 4 (**Fig. 2A**). Both SBP and DBP in SHRSP rats were markedly significantly than for the control rats. At month 14, SHRSPs exhibited prone position without movement, inability to walk, and depression-like behavior. Histopathological examination was performed on the brains and kidneys of three rats. As shown in **Figure 2B**, Rat 1 showed no observable changes in the brain, with only minimal swelling, degeneration, and fibrosis in some of the glomeruli and tubules. In Rat 2, hemorrhagic foci were visible around the hippocampus and ventricles, with necrosis and atrophy in the glomeruli and tubules, some glomerular hyaline degeneration and fibrosis, thickened arteriole walls, and inflammatory cell infiltration. In Rat 3, multiple scattered hemorrhagic foci were found in the cortical and hippocampal regions of the brain, with necrosis and atrophy in the glomeruli and tubules, some glomerular hyaline degeneration and fibrosis, thickened arteriolar walls, and multiple infiltrating inflammatory cells around narrowed lumens. The SHRSP rats exhibited clinical characteristics of elevated blood pressure, along with varying degrees of cerebral lesions and renal pathology. Overall, the SHRSP rat model mimics the onset and progression of hypertension.

**Fig. 2.**
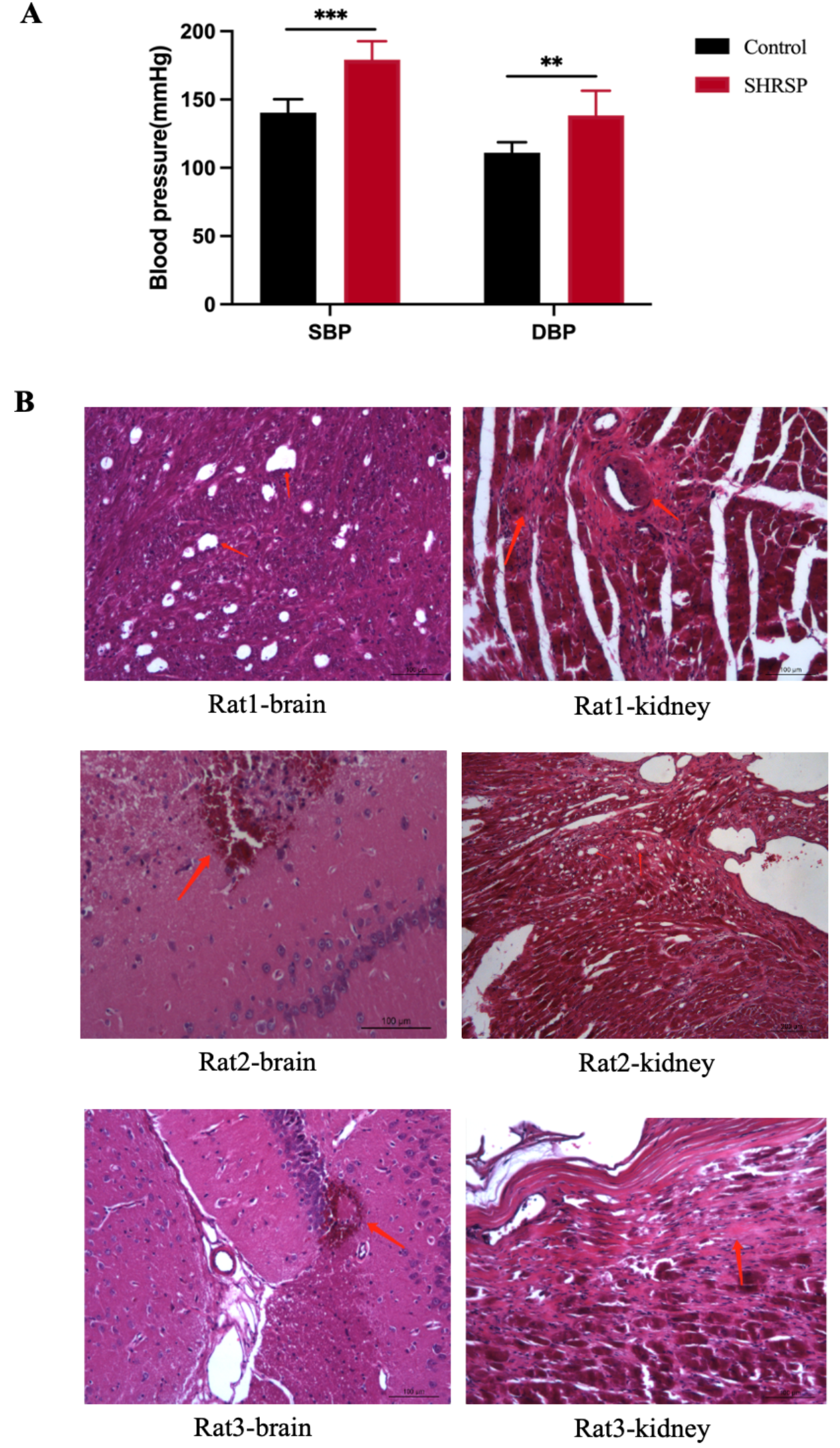
Blood pressure and histopathology in SHRSP rat model. (A) Blood pressure of SHRSP rats at 4 months, SHRSP represents experimental group (n=18), Control represents control group (n=6), SBP is systolic blood pressure, DBP is diastolic blood pressure, ** P < 0.01,*** P < 0.01; (B) Brain and renal histopathology of SHRSP rats at 14 months, HE staining, 20 times magnification.

### Urinary proteome changes in SHRSP rats

To investigate the urine proteome changes in the progression with hypertension, urine samples from four SHRSP rats and from six time points (months 1, 4, 8, 10, 12, and 14) were analyzed by label-free DIA-MS/MS quantitation. A total of 884 urinary proteins were identified with FDR < 0.01. After the missing values of proteomic data was imputed with the sequential-KNN method, a total of 628 proteins were retained for subsequent differential urinary protein selection. Considering the complexity of hypertension and that disease progression and clinical manifestations of individuals might different, the self-comparison approach was adopted to screen for differential proteins. The screening criteria was as follows: fold change ≥ 2 or ≤ 0.5, P < 0.05. For Rat1, a total of 173, 321, 91, 61 and 52 differential proteins were significantly changed at Month 4, 8, 10, 12, and 14, respectively (**Fig.3A, Table S1**). The overlap of these differential proteins screened at five time points is presented in **Fig. 3B**, only one protein was identified at these time points in common. For Rat2, a total of 176, 302, 128, 113 and 65 differential proteins were significantly changed at Month 4, 8, 10, 12, and 14, respectively (**Fig.3A, Table S1**). The overlap of these differential proteins screened at five time points is presented in **Fig. 3B**, only two protein was identified at these time points in common. For Rat3, a total of 113, 380, 98, 69 and 41 differential proteins were significantly changed at Month 4, 8, 10, 12, and 14, respectively (**Fig.3A, Table S1**). The overlap of these differential proteins screened at five time points is presented in **Fig. 3B**, only one protein was identified at these time points in common. For Rat4, a total of 126, 214, 261, 29 and 95 differential proteins were significantly changed at Month 4, 8, 10, 12, and 14, respectively (**Fig. 3A, Table S1**). The overlap of these differential proteins screened at five time points is presented in **Fig. 3B**, only two protein was identified at these time points in common. The above results showed that most of differential proteins changed uniquely at multiple different time points, indicating that different biological changes may occur during the progression of hypertension.

**Fig. 3.**
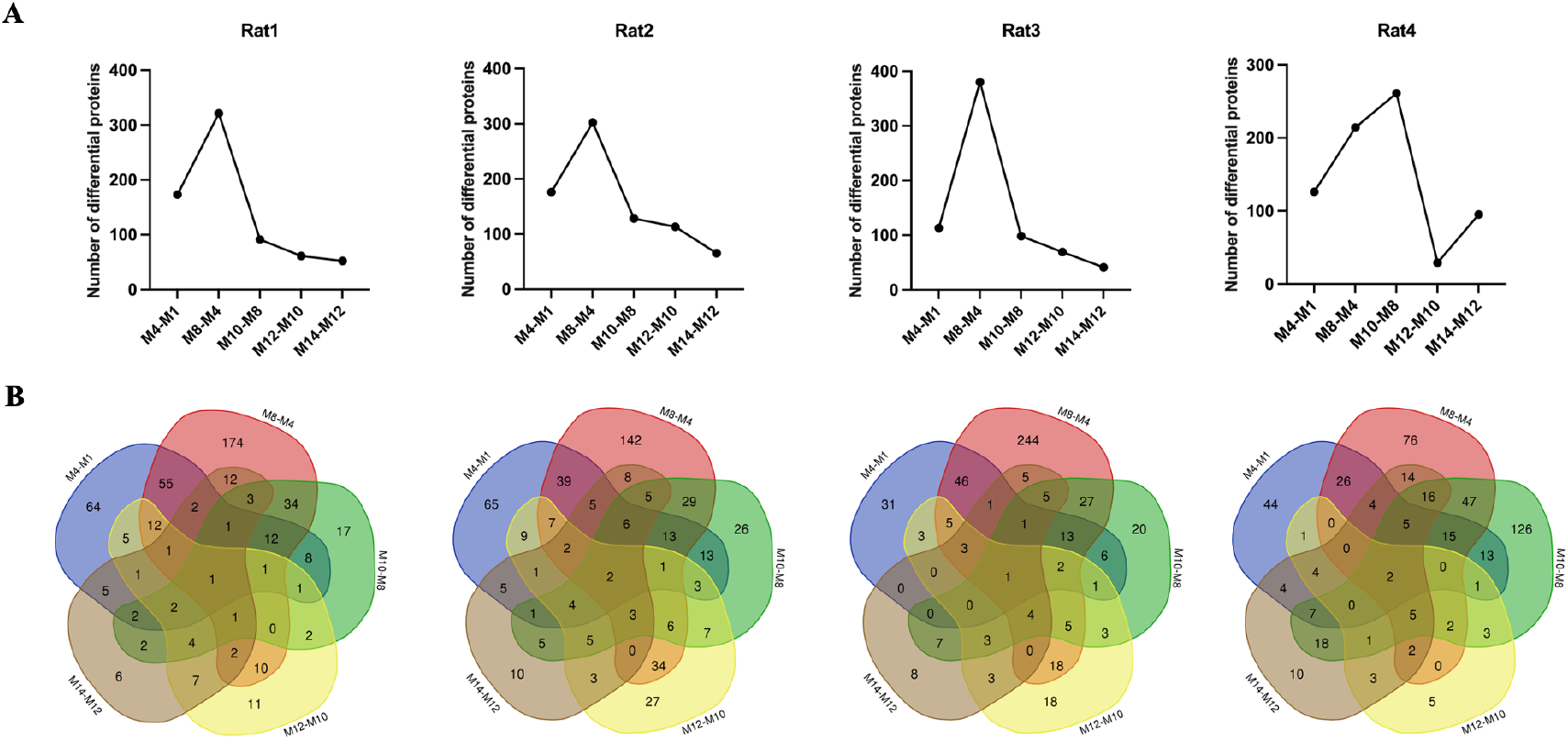
Proteomics changes of urine samples in SHRSP rat model. (A) Changes of number of differential urinary proteins identified at multiple time points in SHRSP rats. (B) The Venn diagram of the differential urinary proteins identified at multiple time points in SHRSP rats.

### Functional analysis of differential in SHRSP rats

To explore the changes of pathophysiological functions at different stages in the progression of hypertension, functional annotation was performed on the differential proteins identified at months 4, 8 and 14, including the biological processes and pathways changes.

On Rat1, in the biological process category (**Fig 4A**), cellular oxidant detoxification and proteolysis were mainly enriched at all time points, glycolytic process was only enriched at months 4; blood coagulation and inflammatory response were only enriched at months 8; regulation of systemic arterial blood pressure by renin-angiotensin was only enriched at months 14. In the pathway category (**Fig 5A**), apelin adipocyte signaling pathway, iron homeostasis signaling pathway and NRF2-mediated oxidative stress response were enriched at months 4; coagulation system and calcium transport I were enriched at months 8; glucocorticoid receptor signaling was enriched at months 14.

**Fig. 4.**
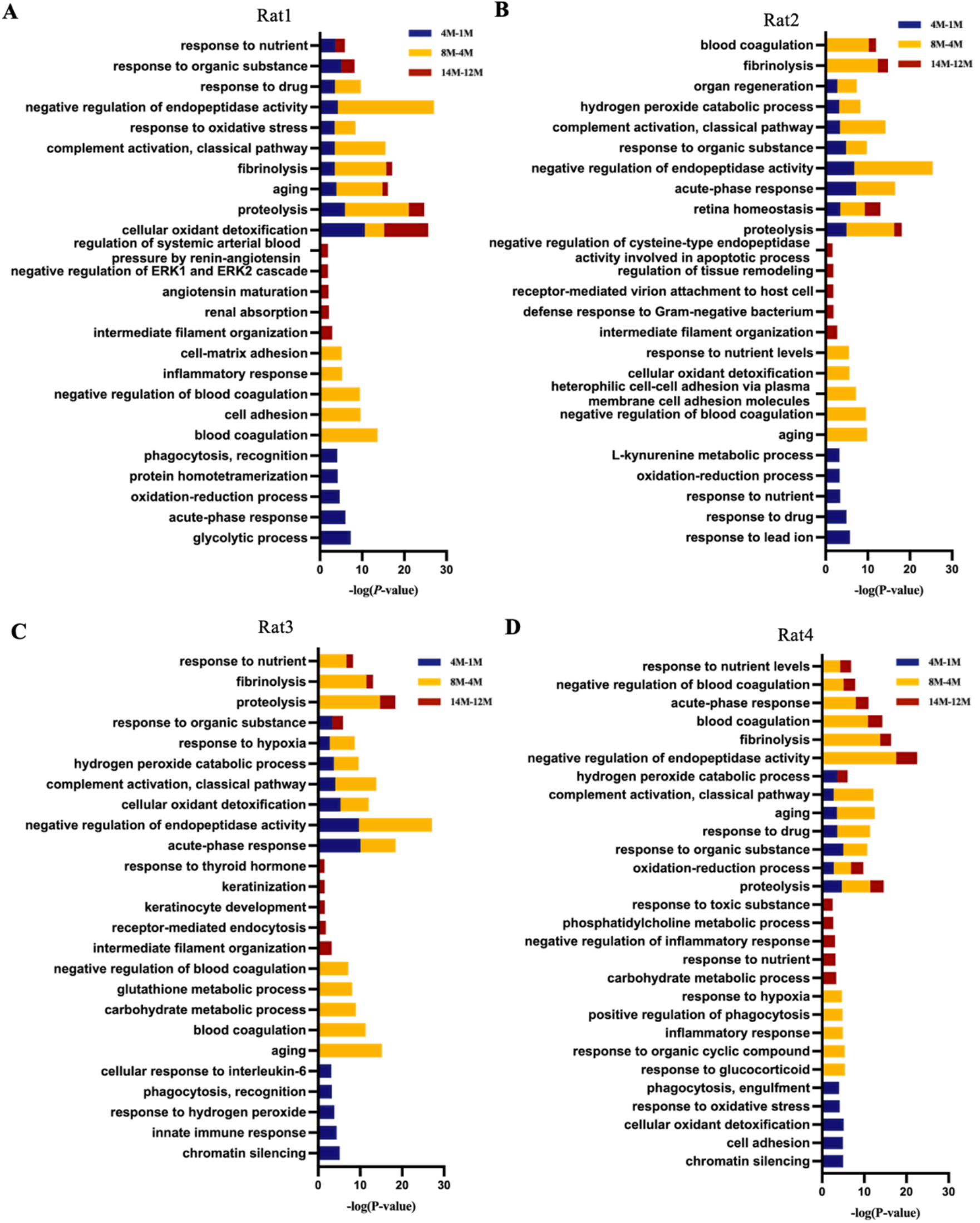
Biological processes enrichment analysis of differential proteins in the progression of hypertension in SHRSP rats. The top 5 biological processes of differential proteins at multiple time points in SHRSP rats. All enrichment items were consistent with P < 0.05 standard.

**Fig. 5.**
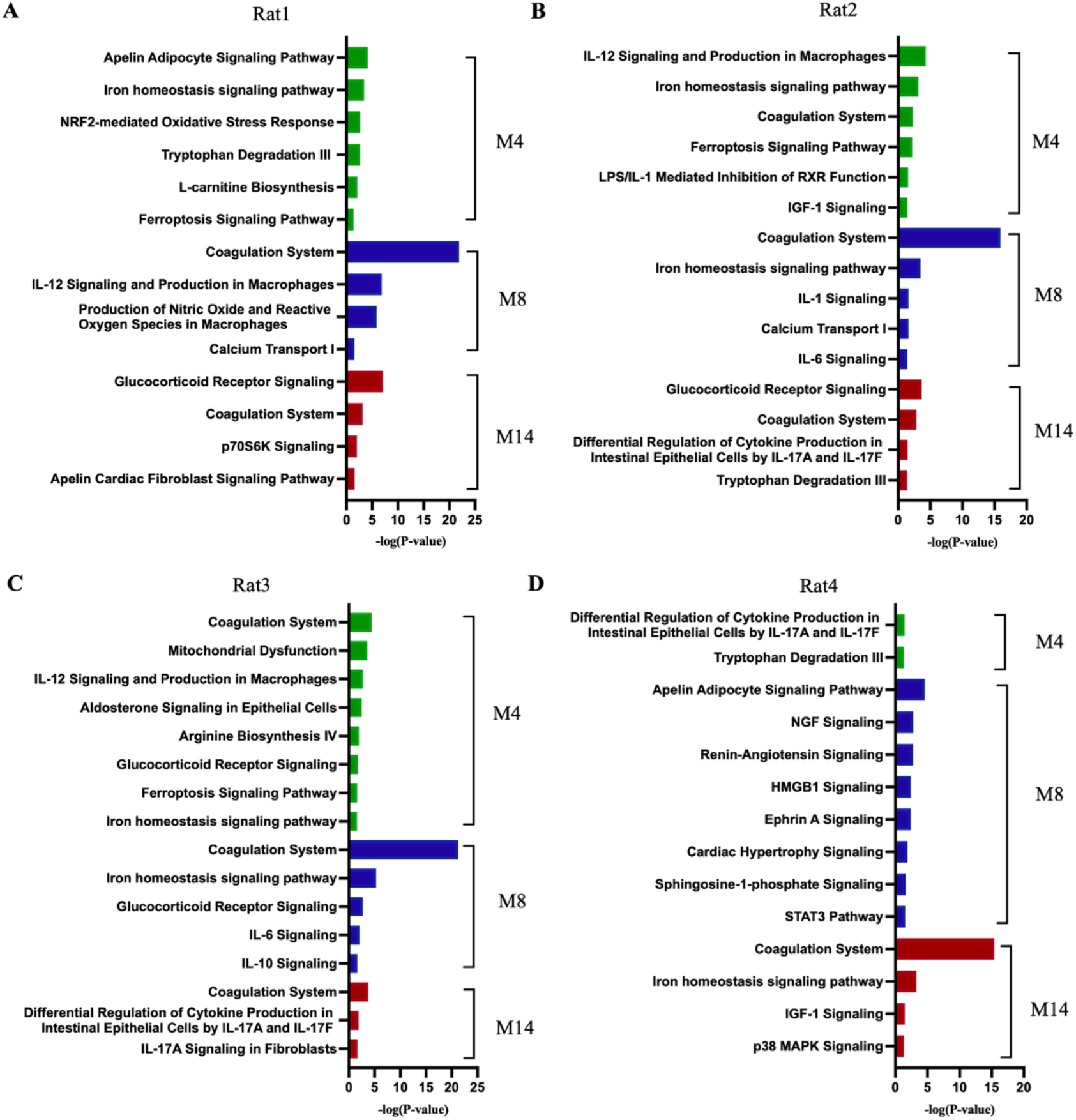
Canonical pathway enrichment analysis of differential proteins in the progression of hypertension in SHRSP rats. The representative pathways of differential proteins at multiple time points in SHRSP rats. All enrichment items were consistent with P < 0.05 standard.

On Rat2, in the biological process category (**Fig 4B**), proteolysis and retina homeostasis were mainly enriched at all time points, response to lead ion was only enriched at months 4; negative regulation of blood coagulation was only enriched at months 8; intermediate filament organization and defense response to Gram-negative bacterium were only enriched at months 14. In the pathway category (**Fig 5B**), IL-12 Signaling and production in macrophages and IGF-1 signaling were enriched at months 4; IL-1 signaling, calcium transport I and IL-6 signaling were enriched at months 8; glucocorticoid receptor signaling was enriched at months 14.

On Rat3, in the biological process category (**Fig 4C**), acute-phase response and complement activation classical pathway were enriched at months 4 and 8, chromatin silencing and innate immune response were only enriched at months 4; aging and blood coagulation were only enriched at months 8; intermediate filament organization was only enriched at months 14. In the pathway category (**Fig 5C**), coagulation system was enriched at all time points; glucocorticoid receptor signaling was enriched at months 4 and 8; IL-10 signaling was enriched at months 8; IL-17A signaling in fibroblasts was enriched at months 14.

On Rat4, in the biological process category (**Fig 4D**), proteolysis and oxidation-reduction process were enriched at all time points; chromatin silencing and cell adhesion were only enriched at months 4; response to glucocorticoid was only enriched at months 8; carbohydrate metabolic process was only enriched at months 14. In the pathway category (**Fig 5D**), tryptophan degradation III was enriched at months 4; renin-angiotensin signaling and STAT3 pathway were enriched at months 8; p38 MAPK signaling was enriched at months 14.

Overall, the above results showed that the changes of hypertension can be reflected in urine proteins and the biological processes and pathways enriched at the same time point in the progression of hypertension differed significantly among different rat individuals.

## Discussion

Hypertension is a multifactorial disease involving environmental and genetic factors as well as risky behaviors,has become the most critical and expensive public health problem. The pathophysiological mechanisms of hypertension are still not well understood. In this study, we systematically investigated dynamic changes in urinary proteome in the SHRSP rats during hypertension development.

We found that some important biological functions or pathways enriched in the SHRSP rats were associated with the pathological mechanism or drug targets of hypertension. For example, i) regulation of systemic arterial blood pressure by renin-angiotensin and renin-angiotensin signaling were enriched in the SHRSP rats. Systemic arterial hypertension is characterized by persistently high BP in the systemic arteries(28). Blockers of the renin-angiotensin-aldosterone system (RAAS) is the cornerstone in the treatment of hypertension(29). ii) Response to glucocorticoid and glucocorticoid receptor signaling were enriched in the SHRSP rats. Glucocorticoids participate in regulating blood pressure through various extrarenal tissues. Excess glucocorticoid levels cause promiscuous activation of mineralocorticoid receptors and induce hypertension. Glucocorticoid receptors are widely expressed in many organ systems involved in blood pressure regulation and play an important role in the pathogenesis and maintenance of hypertension(30). iii) Calcium Transport I was enriched in SHRSP rats. Calcium channel blockers (CCBs) inhibit the flow of extracellular calcium through ion-specific channels across the cell membrane. Although several types of calcium channels have been identified, currently available CCBs inhibit L-type channels in humans. When inward calcium flow is inhibited, vascular smooth muscle cells relax, leading to vasodilation and lower BP (31). iv) Aldosterone adipocyte signaling pathway was enriched in SHRSP rats. Aldosterone is a steroid hormone that is synthesized and secreted by the glomerular zona in the outer layer of the adrenal cortex, regulates sodium homeostasis, and controls blood volume and blood pressure. Excess secretion of this hormone may lead to hypertension and exacerbate disease morbidity and mortality. An understanding of the signaling pathway that regulates aldosterone biosynthesis may help researchers identify new targets for therapeutic intervention in cardiovascular diseases such as hypertension(32). v) Apelin adipocyte signaling pathway was enriched in SHRSP rats. Apelin is a vasoactive peptide and its receptor APJ are widely expressed in the cardiovascular regulatory regions of blood vessels, heart and brain in the cardiovascular system. Based on accumulating evidence, the apelin/APJ receptor system plays a regulatory role in cardiovascular physiology and pathophysiology, making it a potential target for cardiovascular drug discovery and development (33). vi) Oxidative stress response was enriched in SHRSP rats. Studies have shown that oxidative stress is involved in the pathogenesis of hypertension (34), and oxidative stress promotes endothelial dysfunction, vascular remodeling and inflammation in the pathophysiological process of hypertension, leading to vascular damage (35). Clinical studies of patients with essential hypertension have shown that blood pressure is positively correlated with the levels of oxidative stress biomarkers and negatively correlated with antioxidant levels (36). Urinary proteomics will increase our understanding of the pathogenesis and progression of hypertension and will someday be used to monitor the progression of hypertensive.

The duration of hypertension development and its relevance to age is a challenge when conducting long-term longitudinal studies in humans. It is important to note that disease progression may differ among individual animals, so proteomic analyses of multifactor complex diseases can be performed in ways that compare individual animals. At the same time, in order to reduce the impact on growth and development, this study conducted differential protein screening by comparing the time points before and after the occurrence of hypertension in a single rat. We found several important functional annotations were identified in SHRSP rats, suggesting that a limited number of important pathways may be involved in hypertension. In addition, we found that although they are all hypertensive rats, different individuals may be involved in different pathological mechanisms in the occurrence and development of hypertension. The unique biological pathways of different rats may provide new insights into the personalized diagnosis and treatment of hypertension. In the future, the pathway analysis can be conducted in patients with hypertension by analyzing the urinary proteome, and targeted drugs may be selected for personalized treatment.

## Conclusions

This study was a preliminary study with a small sample size. These results showed that the changes of hypertension can be reflected in urine proteins. The urine proteomics has the potential to be used to study the mechanisms of hypertension, discovery new drug targets, and provide personalized antihypertensive treatment strategies.

## Animal Ethics

The following information was supplied relating to ethical approvals:

The experiment was approved by Peking Union Medical College (Approval ID: ACUC-A02-2014-007).

## Data availability

Data will be made available on request.

